# Early Life Adversity, but not suicide, is associated with less prefrontal cortex gray matter in adulthood

**DOI:** 10.1101/531871

**Authors:** Mark D. Underwood, Mihran J. Bakalian, Teresa Escobar, Suham Kassir, J. John Mann, Victoria Arango

## Abstract

**Background:** Suicide and major depression (MDD) are more prevalent in individuals reporting early life adversity (ELA). Prefrontal cortex volume is reduced by stress acutely and progressively *in vivo*, and changes in neuron and glia density are reported in depressed suicide decedents. We previously found reduced levels of the neurotrophic factor BDNF in suicide decedents and with ELA, and in the present study we sought to determine whether cortex thickness, neuron density or glia density in the dorsolateral prefrontal (BA9) and anterior cingulate (BA24) cortex are associated with ELA or suicide.

**Methods:** A total of 52 brains, constituting 13 quadruplets of nonpsychiatric nonsuicide controls and MDD suicide decedents with and without ELA (n=13/group), all with psychological autopsy, were matched for age, sex and postmortem interval. Brains were collected at autopsy and frozen and blocks containing BA9 and BA24 were later dissected, post-fixed and sectioned. Sections were immunostained for NeuN to label neurons and counterstained with thionin to stain glial cell nuclei. Cortex thickness, neuron and glial density and neuron volume were measured by stereology.

**Results:** Cortical thickness was 6% less with an ELA history in BA9 and 12% less in BA24 (*p*<0.05), but not in depressed suicide decedents in either BA9 or BA24. Neuron density was not different in ELA or in suicide decedents, but glial density was 17% greater with ELA history in BA9 and 15% greater in BA24, but not in suicides. Neuron volume was not different with ELA or suicide.

**Discussion:** Reported ELA, but not the stress associated with suicide, is associated with thinner prefrontal cortex and greater glia density in adulthood. ELA may alter normal neurodevelopment and contribute to suicide risk.

## INTRODUCTION

Stress and mental illness, including major depressive disorder (MDD), are associated with volumetric loss in medial prefrontal cortex and hippocampus *in vivo* (see (Belleau et al., 2018) for a recent review). The stress can be recent and associated with the mood disorder or distant and occurring in childhood, and both types of exposures can be associated with smaller brain regional volume. The early life stress-associated volume loss in the prefrontal cortex and hippocampus is hypothesized to be the result of severe stress impacting neuronal development at a critical period (Crews et al., 2007).

Early life adversity (ELA) is associated with increased risk for suicide (Johnson et al., 2002; Labonte and Turecki, 2012; Turecki et al., 2012; Dykxhoorn et al., 2017; Youssef et al., 2018) and adult depression (Miniati et al., 2010; Harkness et al., 2012; C. Heim and Binder, 2012; Nanni et al., 2012). We found lower neuron density in depressed suicide decedents and hypothesized lower neuron density contributes to higher levels of postsynaptic 5-HT_1A_ and 5-HT_2A_ receptors (M. D. Underwood and Arango, 2011; M.D. Underwood et al., 2018). We also find higher levels of 5-HT_1A_ and 5-HT_2A_ receptor binding associated with early life adversity (M.D. Underwood et al., 2018). Changes in the density of neurons and glia have been reported in MDD by others (Rajkowska et al., 1999; Rajkowska, 2000; Cotter et al., 2001). Suicide and the mood disorder accompanying suicide are sources of current profound stress, potentially resulting in elevated cortisol and glutamate and in reduction in the brain trophic molecule BDNF (Deveci et al., 2007; Kim et al., 2007; Dwivedi, 2012). We found lower levels of BDNF in the anterior cingulate cortex in individuals reporting ELA and in suicide decedents (Youssef et al., 2018) raising the possibility that less BDNF may lead to reduced trophic neuronal support and reduced brain region volume. Therefore, reduced volume of the prefrontal cortex due to ELA could be due to loss of neurons, less interneuronal parenchyma, fewer glia, less vascular tissue or some combination of these effects.

In the present study, we sought to determine whether there is a loss of thickness of prefrontal cortex associated with reported early life adversity (ELA) and to distinguish effects of ELA from those associated with suicide. We also set out to determine whether the thickness of the cerebral cortex is associated with neuron density or glia density and how those associations are similar or different in depressed suicide decedents or nonsuicide controls with or without reported ELA.

## METHODS

The respective Institutional Review Boards for Human Use Considerations of the New York State Psychiatric Institute, the University of Pittsburgh, and the Republic of Macedonia approved the procedures for collection and use of brain tissue.

Brain tissue samples were obtained from the respective medical examiner’s offices from which the decedents originated. Upon removal of the brain from the cranium, the dura mater was stripped, and the brainstem and cerebellum were removed. The brain was bisected and the right hemicerebrum was cut into 2 cm thick sections in the coronal plane. The tissue slabs were placed on a glass plate, frozen in liquid R-12 (Freon 12, dichlorodifluoromethane), placed in pre-labelled plastic bags and kept frozen at −80 °C. A sample of cerebellar tissue was collected and used for brain toxicological analyses. The remaining tissue was placed in formalin for gross and microscopic neuropathological examination.

A total of fifty-two subjects were assigned to quadruplets, consisting of a nonsuicide with no reported history of ELA, a suicide with no reported ELA, a nonsuicide with ELA, and a suicide with ELA, based on age (±5 years), sex, race, and postmortem interval (± 5 hours and < 24 hours). Control subjects without ELA (n=13) died of sudden death (n=9) or natural causes (n=4). Nonsuicides with reported ELA (n=13) died of heart attack (n=10), or accidental sudden death (n=3). Suicides without a reported ELA died from hanging (n=6), gunshot wound (n=2), physical trauma (n=3), or poisoning (n=2). Suicides with a reported ELA died from hanging (n=6), gunshot wound (n=2), suffocation (n=1), or poisoning (n=4). See Table 1 for subject characteristics. All subjects were free of gross neuropathology and had negative toxicology screens in blood, urine, and bile for psychoactive drugs except for two suicide cases which were positive for benzodiazepines, one of whom also positive for barbiturates.

**Table 1.** Group Demographics

| Group | Age<br>(years) | Sex<br>(M:F) | Race<br>(W:B:H) | PMI<br>(hours) | Brain pH | Cause of Death | Axis I Diagnosis (n) |
| --- | --- | --- | --- | --- | --- | --- | --- |
| Nonsuicide,<br>No ELA<br>(n=13) | 37±4 | 12:1 | 12:0:1 | 14.2±1.5 | 6.5±0.11 | Cardiovascular (4),<br>Accident (6), Homicide<br>(3) | None (13) |
| Suicide, No<br>ELA (n=13) | 37±5 | 12:1 | 12:1:0 | 15.1±1.4 | 6.44±0.08 | Hanging (6), GSW (2),<br>Physical Trauma (3),<br>Poisoning (2) | None (2), MDD (9), Schizophrenia (1) |
| Nonsuicide,<br>with ELA<br>(n=13) | 38±5 | 12:1 | 7:4:2 | 13.2±1.1 | 6.56±0.07 | Cardiovascular (10),<br>MVA (1), Accidental<br>Electrocution (1),<br>Hemorrhage (1) | None (11), Gambling Disorder (1),<br>Conduct Disorder (1) |
| Suicide, with<br>ELA (n=13) | 36±5 | 12:1 | 9:1:3 | 19.2±2.4 | 6.58±0.07 | Hanging (6), GSW (2),<br>Suffocation (1),<br>Poisoning (4) | MDD (10), MDD, Gambling Disorder, OCD (1),<br>Conduct Disorder (1), MDD, Conduct<br>Disorder, Cannabis Abuse, ADHD (1) |

**Abbreviations:** ELA: Early Life Adversity, MVA: Motor Vehicle Accident, GSW: Gunshot Wound.
Age, PMI, and Brain pH are expressed as mean ± S.E.M.

### Tissue Preparation

Tissue blocks containing BA9 and BA24 were dissected from frozen coronal slabs of the right hemisphere. The blocks were sectioned 60 μm on a freezing microtome (Microm HM 440E, Walldorf, Germany) and slides were stored until used.

### Immunohistochemistry

The sections on slides (3 sections from each Brodmann area in each ID at 1 mm intervals) were fixed in 4% paraformaldehyde/0.1M phosphate buffer, pH 7.4 for 30 minutes. Then the sections were washed in 50 mM PBS, treated with 0.3% hydrogen peroxide, washed again and blocked for 2 hours in NHS. The sections were incubated overnight in primary anti-NeuN mouse monoclonal antibody (1:20000; Chemicon, Temecula, CA, Figure 1) in 0.1% NHS and 0.3% Tx-100 at room temperature. Adjacent sections at 1 mm intervals were stained with cresyl violet.

**Figure 1.**
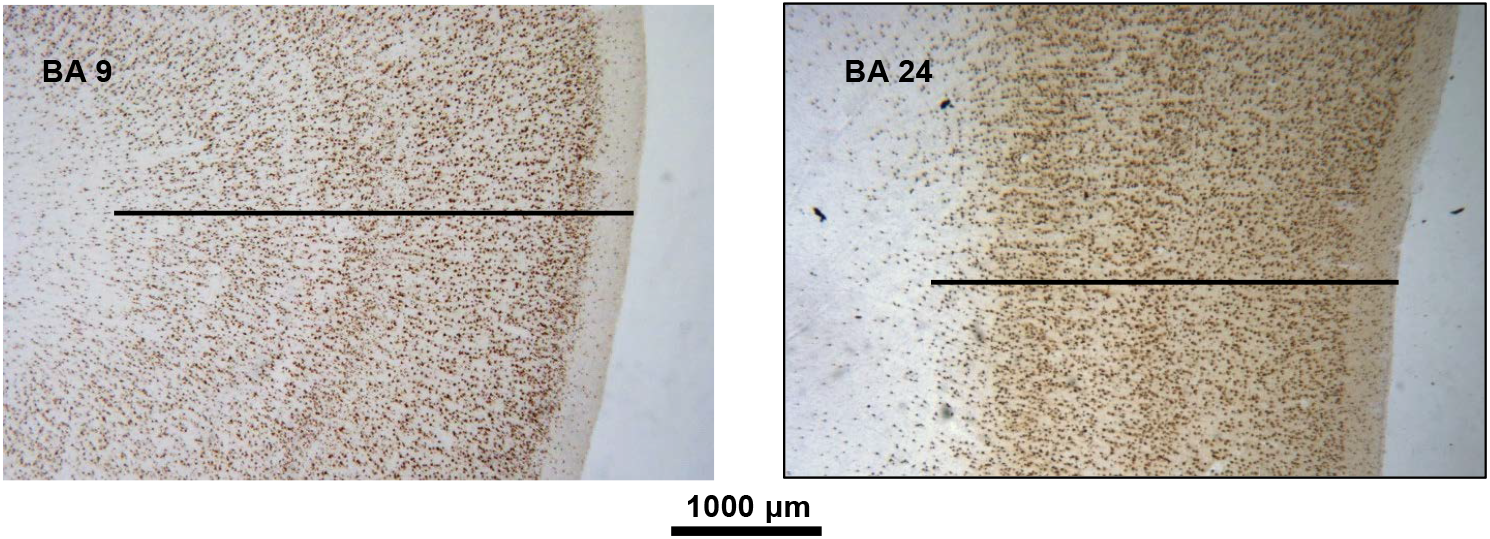
NeuN immunocytochemistry in representative tissue sections in BA9 and BA24. Cortex thickness measurement is depicted by the transept line.

### Stereology

Length distances for each microscope objective were calibrated for linear measure by the software (StereoInvestigator, MBF Bioscience, Williston, VT). The thickness of the cortex was measured at 40X drawing a transept line perpendicular to outer margin of layer I extending to the layer VI-white matter interface (Figure 1).

The total cell number could not be determined since the entirety of BA9 nor BA24 was available for sampling, making NeuN^+^ and glial cell density the best possible metrics. The density of NeuN-IR^+^ neurons in BA9 and BA24 was determined stereologically using a personal computer-based imaging workstation equipped for stereology (StereoInvestigator, MBF Bioscience). The computer is attached to a microscope (Leica, model Diaplan; Wetzlar, Germany) fitted with a motorized stage (Ludl Electronic Products, Hawthorne, NY) and stage position encoder (Heidenhain MT12, Schaumburg IL). Neuronal and glial density was determined by 3-dimensional stereology using the fractionator method (e.g., (Gundersen et al., 1988b; Gundersen et al., 1988a). Slides were initially viewed at 16X total magnification under a Leica Wild M3Z stereoscope and BA9 and BA24 margins were demarcated on each slide of each case using the Brodmann area map of Rajkowska and Goldman-Rakic (Rajkowska and Goldman-Rakic, 1995b, a). The assumption was made that neuron density and glia density were representative of the region of interest throughout the rostrocaudal extent of the respective Brodmann area. We sought to maximize the sampling of each Brodmann area while having an adequately small coefficient of variation and sampled 3 slides 1 mm apart for each Brodmann area in each case. BA contours were drawn by the image analyst with the use of the computer pointing device and the microscope motorized stage. The contour corresponded to the recognized BA boundary. The *disector* began 3 μm below the cut edge of the section. Immuno-positive neuron “tops” and glial cell nuclei were defined at a total magnification of 400X. Neuronal cell volumes were acquired using the nucleator probe (Gundersen, 1988; Gundersen et al., 1988b).

### Statistical Analysis

Statistical tests were done using SPSS (Version 24, IBM Analytics, NY) and R (Version 3.5.1, R Foundation for Statistical Computing; https://cran.r-project.org).

There were three primary hypotheses: 1) cortical thickness is less in suicides; 2) there is a lower density of neurons in the PFC in suicide; and, 3) there is a greater density of glia in the PFC in suicides. Linear models were used since the response variables were continuous (scalar) (SPSS Procedures UNIANOVA, CORRELATIONS, REGRESSION and T-TEST). Suicide and early life adversity were fixed factors, Brodmann area was a random factor. All statistical models included age and sex as covariates, and when found to be significant, correlation analysis was performed. Because there were three hypotheses being tested, Bonferonni correction was employed and p-values were considered statistically significant when p<0.017. Correlations between continuous variables (e.g. thickness correlating with neuronal density or glia density) with age were examined using Pearson Product-Moment Correlation coefficients. Statistical tests were performed on raw values. Cortex thickness, neuron density and glia density were examined in independent tests. Uncorrected p-values are presented in Table 2.

## Results

### Cortical thickness

In non-suicide no ELA controls, cortical thickness in BA9 was 4,630 ± 180 μm and in BA24 was 4,691 ± 252 μm (n=13). The thickness of BA9 (paired t= −1.048; *p*=.3) and BA24 (paired t=0.162; *p*=.872) were not different.

Cortical thickness was not different in suicides in (BA9: F=0.830, df=1,12, *p*=0.380) or BA24 (F=2.861, df=1,12, *p*=0.116). Cortical thickness trended to be less with early life adversity in BA9 (F=5.269, df=1,12, *p*=0.04) and was less in BA24 (F=8.274, df=1,12, *p*=0.014). The effect of ELA was the same in suicides and nonsuicides as there was no interaction between suicide and early life adversity on cortical thickness in either BA9 (F=2.064, *p*=.179) or BA24 (F=0.239, *p*=.638).

### Neuron and glia density

Neuron density in controls (non-suicide and without ELA) in BA9 was 40,611 ± 4090 neurons per mm^3^ and in BA24 was 38,301 ± 2567 per mm^3^ and was not different between regions (t=.750, *p*=.475; Figure 2). Glial density was 58,437 ± 4459 cells per mm3 in controls in BA9 and 58,666 ± 3895 in BA24 and was not different between BA9 and BA24 (Figure 2). Across all groups, glial density (53,471 ± 3696 cells per mm3 in BA9, 53,311 ± 2911 in BA24) was greater than neuron density (44,791 ± 2249 cells per mm3 in BA9, 42,996 ± 1692 in BA24, F=21.779, *p*<0001, repeated measures ANOVA).

**Figure 2.**
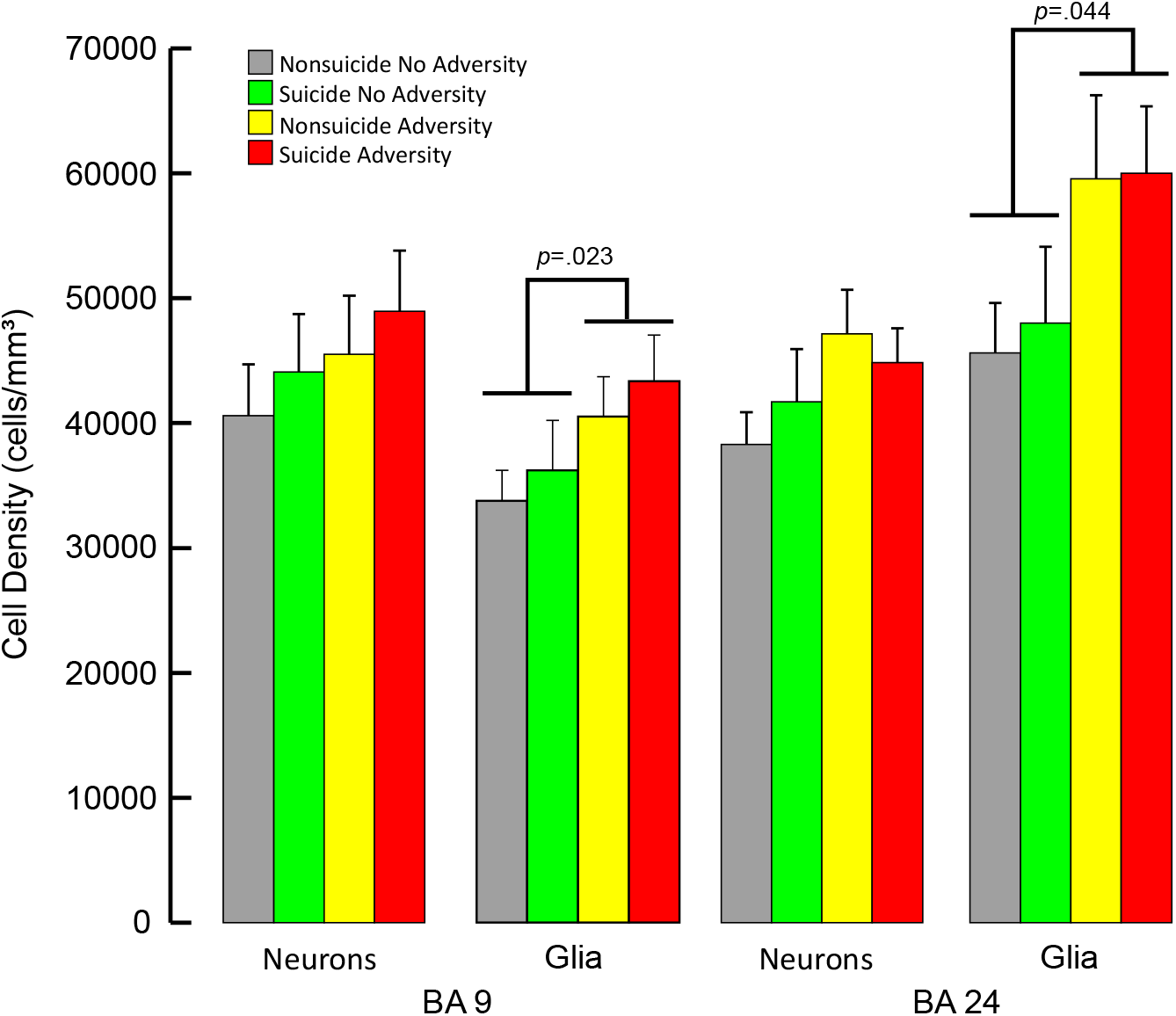
Neuron and Glia Density in BA9 and BA24. In both BA 9 and BA 24, glial density was greater in cases with reported early life adversity, independent of suicide status.

In BA9, neuronal density was not different in suicides (F=1.052, *p*=.329) or in association with early life adversity (F=0.064, *p*=.806). Likewise in BA24, neuron density was also not different in suicide (F=0.212, *p*=.655) or with reported early life adversity (F=1.011, p=.336). Glial density in subjects with early life adversity was greater in BA9 (F=7.813, *p*=.023) and in BA24 (F=5.194, *p*=.044), but the difference was not statistically significant after Bonferonni correction. Glia density was not different in suicide decedents in either BA9 (F=0.025, p=.878) or BA24 (F=0.001, p=0.976). There was no interaction between suicide and early life adversity (F=0.023, *p*=.884). Neuron density correlated negatively with cortical thickness in both BA9 (Pearson coefficient -.338, *p*=.014) but not BA24 (Pearson coefficient .583, *p*=.047).

### Neuron soma volume

Mean neuron soma volume was 2639 ± 94 μm^3^ in BA9 and 2564 ± 80 in BA24 and was not associated with suicide (BA9: F=0.001, *p*=.975; BA24: F=0.282, *p*=.607) or early life adversity (BA9: F=0.242, *p*=.634; BA24: F=2.281, *p*=.159). Similarly, when soma volumes were binned into sizes and compared, the distribution of soma volumes were not different in suicide decedents or with ELA (Figure 3).

**Figure 3.**
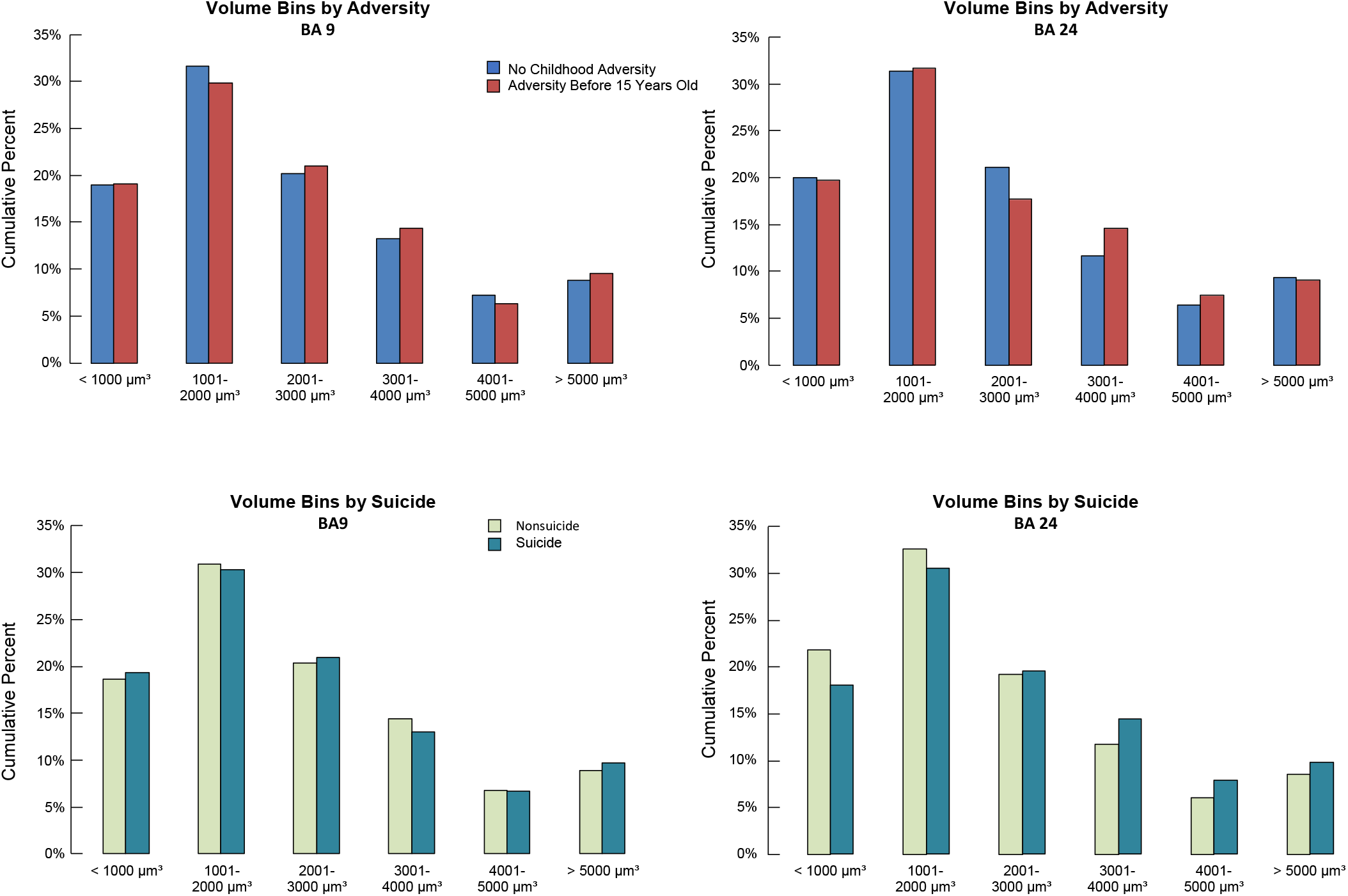
Neuronal cell volume. Neuron soma volume was measured using the Nucleator Method. Soma volumes were aggregated by volume range into bins of 1000 μm^3^. Cell soma volumes did not differ between cases with and without ELA in either BA9 or BA24 (upper panels), nor between nonsuicide and suicide decedents (lower panels)

### Age

Neuron density increases with age in nonsuicides independent of ELA in BA24 (r=.547, *p*=.008). In all cases without ELA, neuronal soma volume decreases with age BA24 (r=-.566, *p*=.006); in that group BA9 thickness also decreases with age (r=-.555, *p*=.006). In cases with ELA, there was no relationship between neuron density, glial density, or neuronal volume with age.

## Discussion

We found that early life adversity but not suicide was associated with less cortical thickness and greater glial density, but no difference in neuron density or neuron soma size.

Cortical thickness is reported to be less in rostral orbitofrontal cortex in MDD postmortem (Rajkowska et al., 1999) and attributed to a shift in the size of neurons from large to small with a decrease in the density of large neurons and an increase in the density of small neurons, suggesting neuronal shrinkage and not neuron loss accounted for the difference in neuron density. Glia are implicated in suicide and the major depression that often accompanies it. Glial density has also been observed to be decreased in density in MDD (Rajkowska et al., 1999; Cotter et al., 2001) in contrast to schizophrenia or Huntington’s disease which are associated with increases in glia density (Rajkowska et al., 1998). Glia include astrocytes, oligodendrocytes and microglia. Astrocytes are specifically implicated in suicide and MDD based on studies using markers specific for astrocytes such as glial fibrillary acidic protein (GFAP). In contrast, studies using nonspecific markers report that astrocytes are increased in density or unchanged (J.J. Miguel-Hidalgo et al., 2000; Davis et al., 2002; Webster et al., 2005). It has been argued that there is a reduction in oligodendrocytes in MDD (Uranova et al., 2004; Vostrikov et al., 2007) that accounts for the overall reduction in glia density in MDD.

Less gray matter volume and cortical thickness are also reported in the frontal cortex in subjects with mood disorders using magnetic resonance imaging by some investigators (see refs (Drevets et al., 1997; Bora and Pantelis, 2011; Belleau et al., 2018)), though not all (Bora and Pantelis, 2011; Gifuni et al., 2016; Rizk et al., 2018). Bora *et al*. (2011) performed a meta-analysis of voxel-based morphometry from 23 studies involving 986 MDD patients and found cortical gray matter volume was reduced in rostral anterior cingulate cortex and dorsolateral and dorsomedial cortex as well as in hippocampus. The smaller gray matter volume was related to illness duration leading the authors to conclude that chronic MDD has deleterious but regionally restricted effects on prefrontal cortex (ibid). The volume deficits related to the MDD, and also to stress (see (Belleau et al., 2018). It is not known, however, whether the reduced cortex thickness were attributable to the mood disorder, a suicide component of the psychopathy, or to both.

The majority of suicide decedents in the current study had MDD. However we did not find cortex thickness different in these MDD suicides in either BA9 or BA24. BA24 was not examined by Rajkowska and colleagues, and 5 of 12 of the MDD cases studied by Rajkowska died from causes other than suicide. Interestingly, the meta-analysis performed by Bora and colleagues (2011) did find smaller BA9 and BA24 gray matter volume in MDD, but there was no indication of whether any of the MDD cases ever made suicide attempts, leaving open the possibility there may be a different effect of suicide and MDD on cortex thickness.

Early life adversity is one of the biggest risk factors for adult suicide and early life adversity can affect the trajectory of neurodevelopment with lasting effects on brain structure and dysfunction (Teicher et al., 2003). Childhood maltreatment can increase the risk of psychiatric problems in adulthood including anxiety, depression and conduct disorder (Gilbert et al., 2009). There is a growing literature in live subjects with early life adversity reporting reduced cortex thickness and or gray matter volume (McCrory et al., 2010). Reduced cortical thickness in maltreated children is reported in the anterior cingulate and superior frontal gyrus (Andersen et al., 2008; Kelly et al., 2013) suggesting an impact on brain morphology from exposure to early life adversity. Such structural differences in the prefrontal cortex and other brain regions may represent an enduring effect on brain development which might lead to the emergence of psychiatric illness later during development or in adulthood (Teicher et al., 2003).

There are critical periods for brain development during which genetic and environmental factors affect neurogenesis, innervation and synaptic pruning and lead to vulnerability for developing psychiatric illness (see (Crews et al., 2007) for review). Andersen and colleagues (2008) found reductions in frontal cortex volume, hippocampus volume, corpus callosum area in women with repeated episodes of childhood sexual abuse. The authors not only concluded there were effects of the childhood sexual abuse on regional brain development, but also that there are sensitive windows during which the brain regions are affected with the frontal cortex most sensitive to abuse at ages 14-16 years. Vulnerability windows may explain why not all individuals exposed to early life adversity have alterations in brain anatomy. Susceptibility to adversity also likely depends on other factors including frequency and intensity of the stress (Edwards et al., 2003), the type of adversity (C. M. Heim et al., 2013; McLaughlin et al., 2014); and genetics (Caspi et al., 2002; Caspi et al., 2003).

The thinner prefrontal cortex and greater glial density associated with early life adversity is suggestive of neuropathology. Acute stress causes alterations in the function of the hypothalamic-pituitary axis, most notably elevated levels of cortisol and cortisol hyperreactivity (Horowitz and Zunszain, 2015). Acute stress elevates glutamate to toxic levels causing dendritic loss (see (McEwen, 2017) for review). Likewise, mood disorders are associated with excess glutamate which can lead to synaptic and neuronal loss (see (Haroon et al., 2017). However, *in vivo* imaging studies have not found associations between cortisol and PFC structure (Drevets, 1999; Gold et al., 2002). Inflammation and neurotoxic sequelae of neuroinflammatory processes are increasingly shown in response to stress and in major depression (Slavich and Irwin, 2014; Setiawan et al., 2015; Lindqvist et al., 2017; van Velzen et al., 2017; Holmes et al., 2018). And higher levels of pro-inflammatory cytokines are associated with thinning of the medial prefrontal cortex (Savitz et al., 2013; van Velzen et al., 2017). We did not find evidence of neuron loss associated with ELA history or death by suicide as there was no difference in neuron density observed in the present study compared to controls. Our measured neuronal density was comparable to those reported in the literature (e.g., (Cotter et al., 2001; M. D. Underwood et al., 2012; Lew et al., 2017), but less than that reported by Rajkowska and colleagues who use a celloidin preparation producing considerable shrinkage and greater cell density (Rajkowska and Goldman-Rakic, 1995a) suggesting that our not finding a difference in neuron density was somehow a confound of the immunostaining or stereologic methods. We did detect a greater density of glial cell nuclei with ELA history and this difference is also not likely a confound since the glia density we measured was comparable to that reported in the literature (e.g., (Öngür et al., 1998; Cotter et al., 2001; J. J. Miguel-Hidalgo et al., 2002).

Serotonergic differences in the prefrontal cortex are reported in depressed suicide decedents (see (M.D. Underwood et al., 2018) for review). Serotonin (5-HT) is also a trophic agent for neurons during neurodevelopment and in adulthood (Booij et al., 2015) and the 5-HT metabolite 5-HIAA is lower in suicide attempters (see (Bach and Arango, 2012; Bach et al., 2013) for recent reviews) raising the possibility that cortical thinning may be the result of neurotransmitter alterations associated with the mental illness. Alternatively, 5-HT is a regulator of BDNF and BDNF has trophic actions on neurons (Mattson et al., 2004). We previously reported lower BDNF levels in anterior cingulate cortex in subjects who had been exposed to ELA and/or died by suicide compared to non-suicide decedents and no reported ELA (Youssef et al., 2018). The 52 cases and controls in the present study were examined for BDNF levels in the study of Youseff et al. (2018). We examined the relationship between BDNF level and the morphometric measures in the present study and found the levels of BDNF correlated with glial density, neuronal volume and cortical thickness (data not shown) raising the possibility that lower amounts of BDNF may underlie the reduced cortical thickness we observed with reported ELA.

### Limitations

Differences in cortex morphometry in depressed suicide decedents do not distinguish between effects attributable to MDD versus those to suicide. Making such a distinction is particularly problematic in postmortem studies when the majority of suicides have MDD or some other mood disorder or psychiatric illness. Suicide has its own associations with cellular changes implicating prefrontal cortex, glia alterations and gray matter volume, that reflect a relationship to the diathesis for suicide, a combination of traits involving mood regulation, decision-making and emotion, social cognitions and learning, that are independent of the associated major psychiatric disorders (for reviews see (Ernst et al., 2009; van Heeringen and Mann, 2014; Balcioglu and Kose, 2018). It is also not possible to determine from postmortem studies, which by their nature examine the brain at a single point in time, whether the thinner cortical thickness reported in studies of MDD patients is state-dependent (and perhaps reversible) or a trait (and indicative of neuropathology), though associations with multiple depression episodes or the summed duration of multiple episodes, suggests the changes in thickness progress with illness duration. Lastly,postmortem studies are not able to demonstrate causality so it is unresolved whether or how the early life adversity leads to increased glia density or reduced cortex thickness.

### Conclusion

Our study went beyond the gross assessment of prefrontal cortex thickness and sought to determine the underlying components of neuron density, neuron size and glial density in neocortex and to separate the effects of depression and suicide from those of early life adversity. We found no differences in neuron density or neuron size in the depressed suicide decedents or with early life adversity, yet cortex thickness was less with early life adversity, suggesting there was neuron loss. Neuron loss, or fewer neurons, with early adversity and combined with the greater glial density may be part of an ongoing pathology or a chronic change. The lack of differences in neuron or glia density in depressed suicides, who suffer severe stress, suggest that acute stress is not as important as childhood adversity or chronic stress experienced over many years. Future studies should not only seek to replicate the finding in a larger sample size but also examine the phenotype of glia involved.

## Acknowledgements

This work was supported by grants from the National Institute of Mental Health (MH40210, MH62185) and the Diane Goldberg Foundation. Some of the brain samples and their psychiatric characterization and storage were funded by NIH grants MH90964 and MH64168. Dr. Mann receives royalties for the commercial use of the C-SSRS from the Research Foundation for Mental Hygiene. Drs. Arango, Underwood and Mrs. Kassir and Mr. Bakalian declare no conflicts of interest.

